# Short-chain PFAS exposure alters embryonic development and behavior in zebrafish

**DOI:** 10.64898/2026.03.03.709373

**Authors:** Zainab Afzal, Evan E Pittman, Vandana Veershetty, Charles Hatcher, Morgan Bailey, Deepak Kumar

## Abstract

Per- and polyfluoroalkyl substances (PFAS) are manmade chemicals that are persistent in the environment and have been linked to various physiological and neurobehavioral outcomes, including anxiety disorders. Trifluoroacetic acid (TFA), a short chain PFAS and the most common PFAS degradation product, is increasingly detected in water, soil, and human blood, raising significant concerns about its developmental toxicity. However, the impact of early-life TFA exposure on neurodevelopment and behavior remain insufficiently characterized. In this study, we employed Zebrafish (*Danio rerio)* embryos as a New Approach Methodology (NAM), to evaluate the development, behavior, and protein expression changes in response to early-life TFA exposure. Embryos were exposed to environmentally relevant low and high concentrations of TFA beginning at one-cell stage. Early developmental physiology was assessed by measuring viability, tail twitch response, hatching rates, and chorion diameters during embryogenesis. Anxiety-like behaviors were evaluated at 5- and 6-days post-fertilization using validated behavioral assays such as the Light-Dark Test and Startle Response. Each test evaluates distinct anxiety-related behaviors by measuring locomotor activity, thigmotaxis (wall preference), and stimulus reactivity, with anxious zebrafish larvae showing increased movement in light and greater wall preference. Then to identify molecular pathways underlying observed developmental phenotypes with TFA exposure, proteomic analyses were performed on embryos at 24- and 48-hours post-fertilization. Our results indicate that TFA exposure altered developmental physiology, evidenced by reduced chorion diameters, and lead to increased anxiety-like behaviors with larvae exhibiting thigmotaxis. These phenotypic changes were accompanied by detectable alterations in the embryonic proteome. Collectively, our findings provide insight into how short-chain PFAS exposure during critical windows of development may contribute to neurobehavioral dysfunction, highlighting potential risks relevant to inform public health policies and environmental regulations.

## Introduction

Per- and Polyfluoroalkyl substances (PFAS) comprise a large class of over 12,000 synthetic chemicals that are widely used in industrial and consumer products due to their oil and water repellent properties. These compounds are composed of strong carbon and fluorine bonds which makes them extremely difficult to breakdown and earning them the designation of “forever chemicals.” As a result, PFAS are highly resistant to environment degradation and consequently bioaccumulate in humans and wildlife. PFAS contamination has been detected across multiple environmental and biological samples, including water, soil, blood, and serum (Ahrens & Bundschuh, 2014; Fenton et al., 2021; Quinete, 2009). Epidemiological studies have linked PFAS exposure to a wide range of adverse health outcomes including metabolic disruption (Gaillard et al., 2025), immune dysfunction (Martano et al., 2025), and neurobehavioral alterations (Kebieche et al., 2025). Notably, elevated PFAS exposure in humans has been associated with increased anxiety-like behavior in adults (Hua et al., 2025), while prenatal and early life exposure for infants has been linked to impaired motor development and decreased lean mass (Van Beijsterveldt, 2025; Varsi, 2022).

Thus far, there have been regulatory and toxicological efforts that have focused on long-chain PFAS, such as perfluorooctanoic acid (PFOA). However, increasing restrictions on these compounds have driven a transition toward short-chain PFAS, including trifluoroacetic acid (TFA), under the assumption that they represent a safer alternative (Brendel, 2018). With this assumption TFA is not currently strictly regulated, and is increasingly being detected in water, soil, and human blood (Zhai, 2015; Zhi, 2024). Although TFA is generally considered less toxic than long-chain PFAS, on a per-molecule basis, the rising pollutant levels of TFA are over double or triple the amount of PFOA, which in essence can cause TFA to have more toxic effect due to the increased levels of exposure (Arp, 2024; Sodani, 2025; Zheng et al., 2024). Additionally, as TFA is a common degradation product, historical human blood serum samples tested for presence of PFAS, have also detected high levels of TFA (Cheng et al., 2025). Another short-chain PFAS, bis(trifluoromethane)sulfonimide lithium salt (HQ115), used in electric vehicle batteries, has been shown to induce kidney injury and disrupt genes involved in inflammation and uric acid metabolism (Sands et al., 2024). It is also increasingly being detected in surface and wastewater near certain manufacturing sites (EPA, 2023). Despite these trends, relatively few studies have examined the developmental and neurobehavioral impacts of such short-chain PFAS, particularly during early life stages.

We employed the zebrafish (*Danio rerio*), a well-established vertebrate model species whose embryos serve as a New Approach Methodology (NAM) (Ball et al., 2025; Kalueff, 2014), and as a powerful system to investigate TFA’s toxic effects during development. Zebrafish embryos develop externally and have clear chorions, which allows direct observation of their early developmental processes. Their small size and high progeny counts with synchronized development allows for multiple molecular and behavioral tests to be performed in a shorter amount of time. Additionally, zebrafish is a vertebrate system which shares about 70% of its genome with humans, allowing for direct comparisons of genes and proteins to gain mechanistic insights into human diseases (Howe, 2013).

In this study, we used zebrafish embryos and larvae to examine the developmental effects, behavioral outcomes, and proteomic changes in response to TFA exposure. Specifically, we assessed early developmental points, at 24 hours post-fertilization (hpf) when organogenesis in underway (Kimmel et al., 1995), to assess for changes in chorion sizes and stimulus evoked tail twitching, and used later 6 day old larval stage to evaluate for anxiety-like behaviors. We furthered determined whether these behavioral outcomes were associated with changes in protein expressions at 24- and 48-hpf. Collectively, our integrated experimental approach provides a comprehensive evaluation of TFA’s developmental toxicity and challenges the current assumption that TFA is a safer alternative.

## Methods

### Zebrafish husbandry

Adult AB wildtype zebrafish, housed in the Animal Resource Complex on a 14:10 Light:Dark cycle, are maintained in accordance with an approved IACUC protocol # DK06012022. Adults were paired the afternoon, the day before breeding and housed in breeding tanks in a 2:1 female to male ratio. They males and females were separated by a clear central divider along with a mesh at the bottom of the tank to prevent embryo predation. The following morning, the dividers were removed, and the fish were allowed to breed for 15 minutes to obtain developmentally synchronized embryos. After 15 mins, the fertilized embryos were collected and sorted into different control and treatment groups. The embryos were stored in a 28.5°C incubator and the embryo media was changed daily. The embryo media was made RO water buffered with Instant Ocean© sea salt an pH was maintained between 7.0 and 7.5.

### Chemical Preparation and Exposure

Trifluoroacetic acid (TFA; Thermo Scientific, Catalog No. 28901) stock solutions (1 µg/L) were prepared using nuclease-free water. Embryos were housed in 60 mm Petri dishes containing 25 mL of embryo media. Two TFA exposure concentrations were used: a low concentration of 45 ng/L and a high concentration of 1000 ng/L. Concentrations were selected based on reported environmental levels of TFA in rainfall (Wang, 2014).

A 1 µg/L stock solution of HQ115 (Thermo Scientific, Catalog No. 381030500) was prepared in nuclease-free water. Embryos were maintained in 60 mm Petri dishes containing 25 mL of embryo media. Two HQ115 exposure concentrations were used: 400 ng/L (low) and 8000 ng/L (high), based on previously reported dose–response studies (Owens et al., 2023; Sands et al., 2024).

Perfluorooctanoic acid (PFOA; Sigma, Catalog No. 171468-5G) stock solutions (1 µg/L) were prepared in nuclease-free water. Exposure conditions matched those described above, with embryos maintained in 60 mm Petri dishes containing 25 mL of embryo media. Two PFOA concentrations were used: 700 ng/L (low) and 10,400,000 ng/L (high). Concentrations were determined based on water level regulations and presence in serum levels of children (Cordner et al., 2019; Stein et al., 2013).

### Developmental Assays

Embryos from all treatment groups were assessed at 1- and 24-hours post-fertilization (hpf) to measure chorion diameters. Each treatment group consisted of 50 embryos to account for early developmental mortality. Embryos were imaged using an Olympus MVX10 microscope equipped with CellSens Standard software. Chorion diameter was calibrated with a standardized 1mm scale and quantified using ImageJ.

At 24 hpf, a tail twitch assay was conducted to assess neuromotor responsiveness. Individual embryos were gently stimulated with forceps, and the presence or absence of a tail twitch response to stimuli was recorded manually through visualizations from a dissecting microscope, Ziess SMZ1500. Measurements were performed by multiple independent users, and responses were aggregated to minimize variability arising from differences in applied mechanical force.

### Behavioral Assay

Larvae were transferred daily into fresh embryo media or embryo media and specific PFAS (TFA, HQ115, or PFOA) concentration, depending on the treatment group, until 5 days post-fertilization (dpf). At 5 dpf, larvae were individually placed to a 24-well plate, with one larva per well, for a total of 24 larvae per treatment group. Each well contained 1 mL of embryo media corresponding to the assigned treatment condition. The well plates were then placed into the DanioVision chamber.

Behavioral assays were conducted using the DanioVision chamber equipped with EthoVision software. Using the Ethovision software, behavioral test were designed with an identical setup across all conditions and conducted as a continuous 4 minute assay comprising of alternating light and startle stimulus conditions: minute 1 (light off), minute 2 (light on), minute 3 (light off), minute 4 (mechanical startle stimulus, set to level 4, delivered to plate via tapping). Further, for analysis each well is subdivided into two zone, a center zone (Zone 1) and a periphery zone (Zone 2). Behavioral metrics such as total distance moved and spatial heatmaps are obtained after all conditions are tested. To assess for the persistence of these behavioral effects, the same larvae were retested at 6 dpf using the same protocol.

### Data Analysis

All quantitative data collected from the developmental and behavioral assays were exported into .csv files for analysis. Mean values and statistical significance were calculated between exposure groups and control using students t-tests. Statistical significance was labeled as * if p-value was < 0.05, as ** if p-value was < 0.01, as *** if p-value was < 0.001. All data visualizations were generated using GraphPad Prism.

### Bottom-up Proteomic Analysis

Embryos were collected at 24 hpf and 48 hpf and 25 embryos from each condition were placed in 1.5 ml tubes using a disposable pipette three times. The excess embryo media was removed, and the embryos were flash frozen in liquid nitrogen. Triplicate zebrafish embryos dosed with 0 μM TFA (control), 1.69 nM (low dose), or 25 μM (high dose) at 24- and 48-hours were first ground up using a Spex SamplePrep 2010 Geno/Grinder at a rate of 1000 for two 30 second cycles. The ground embryos were solubilized with 200 μL of 1x PBS buffer and centrifuged at 3,000 rcf for 10 minutes to remove any insoluble debris. The total protein concentration from each sample was measured with a Bradford assay. The protein concentrations ranged from 2.8 mg/mL to 4.5 mg/mL for the 24- and 48-hour TFA samples.

The control, low, and high dose proteins were paired according to their respective exposure time (i.e. 24 or 48 hours). The total protein amount for each of the 3 dosage conditions was normalized to 1 mg/mL for both the 24- and 48-hour timepoints. In total there were 3 samples for the 24-hour control, 3 samples for the 24-hour low dose, and 3 samples for the 24-hour high dose, with an identical setup for the 48-hour samples. The proteins were prepared for LC-MS/MS analysis using a conventional quantitative bottom-up proteomics analysis using 2 isobaric tandem mass tag labeling sets (TMT 10-plex) and the iFASP protocol.^1^ In summary, 200 μL of each 24-hour control and 24-hour dosed protein samples and 150 μL of each 48-hour control and 48-hour dosed protein samples were aliquoted into 8 M urea in 0.1 M Tris-HCl buffer (pH 8.5) to denature the proteins. After performing two buffer exchange steps with the 8 M urea in 0.1 M Tris-HCl the samples were treated with 5 mM TCEP (Tris(2-carboxyethyl)phosphine) and 20 mM MMTS (M-methyl methanethiosulfonate) to reduce and alkylate the disulfide bonds. The protein samples were then labeled using one of the 10 isobaric mass tags from a TMT 10-plex reagent kit following the protocol from Thermo Scientific. The 9 samples from the 24-hour timepoint and the 9 48-hour samples were combined in a way to generate a single 24-hour sample and a single 48-hour sample. Both samples were desalted using Pierce Peptide Desalting columns following Thermo Scientific’s protocol before triplicate LC-MS/MS analyses. The protein samples were analyzed on a Thermo Orbitrap Exploris 480 mass spectrometer coupled to a Thermo Vanquish Horizon LC system. The peptide samples were reconstituted in a 0.1% formic acid in water solution to a final concentration of 1 mg/mL. About 4 μg of total peptide material were injected onto a Thermo Acclaim PepMap C18 1 μm, 100 Å, 1 x 150 mm analytical column. A 115-minute linear gradient of 0 – 40% 0.1% formic acid in 80% acetonitrile at a flowrate of 100 μL/min. The analytical column was heated to a temperature of 50 °C. A cycle time of 1.2 seconds was used to acquire the LC-MS/MS spectra in a data-dependent manner. The resolution and normalized AGC targets of the MS1 and MS2 scans were 120k with 300% and 30k with 100%, respectively. The collision energy was set to 30% with an MS1 scan range of 375 – 1500 m/z and an isolation window of 2 m/z with a dynamic exclusion duration of 30 s. Each peptide sample was analyzed in triplicate by LC-MS/MS.

The LC-MS/MS raw data generated for each timepoint was processed using Proteome Discoverer 3.1 and the SEQUEST HT database search algorithm. The raw spectra were searched against the 2025-08-06 release of the Zebrafish UniProt Knowledgebase. The peptide search parameters included fixed MMTS modifications on cysteine; TMT 10-plex labeling of lysine side chains and peptide N-termini; variable oxidation of methionine; variable deamidation of asparagine and glutamine; and variable acetylation of the protein N-terminus. The digestion enzyme was set as trypsin with a maximum of 2 missed cleavages allowed. There was no normalization mode or scaling applied in the reporter ion quantification settings. Proteins that had a “high” or “medium” FDR confidence (FDR <5%) and did not contain an abundance of 0 in any of the quantification channels were used in subsequent data processing steps. All other parameters were set as default.

The processed data from Proteome Discoverer was normalized by dividing each individual protein abundance by the summed reporter ion intensities of all identified proteins for each sample channel. This accounts for the systematic errors occurring from TMT channel to channel differences (e.g., differential sample loss or TMT labeling efficiency). A fold change ratio was generated between each dose concentration level with the control samples for both the 24- and 48-hour samples. This resulted in 3 ratios for the low vs control and 3 ratios for the high vs control comparisons (in total there are 12 ratios, 6 for each timepoint). The fold-change ratios were log_2_-base transformed, averaged, and subjected to a two-tailed Student’s t-test to generate a p-value for each identified protein. Hit proteins were selected as those with a p-value less than 0.05 and z-score of greater than 1 or less than −1.

## Results

### Early exposure to short-chain PFAS does not impact viability but alters early embryonic development

To assess the impact of early short-chain PFAS exposure on vertebrate development, zebrafish embryos were exposed to trifluoroacetic acid (TFA) via direct addition of different TFA concentrations directly to the embryo media. Embryonic viability was first evaluated to determine whether TFA exposure induced acute developmental toxicity. Exposure to TFA during early development did not significantly affect embryo viability (Figure 1A), with embryo survival being comparable across treatment groups throughout the experimental period, indicating no evidence of acute embryotoxicity with either exposure group.

**Figure 1.**
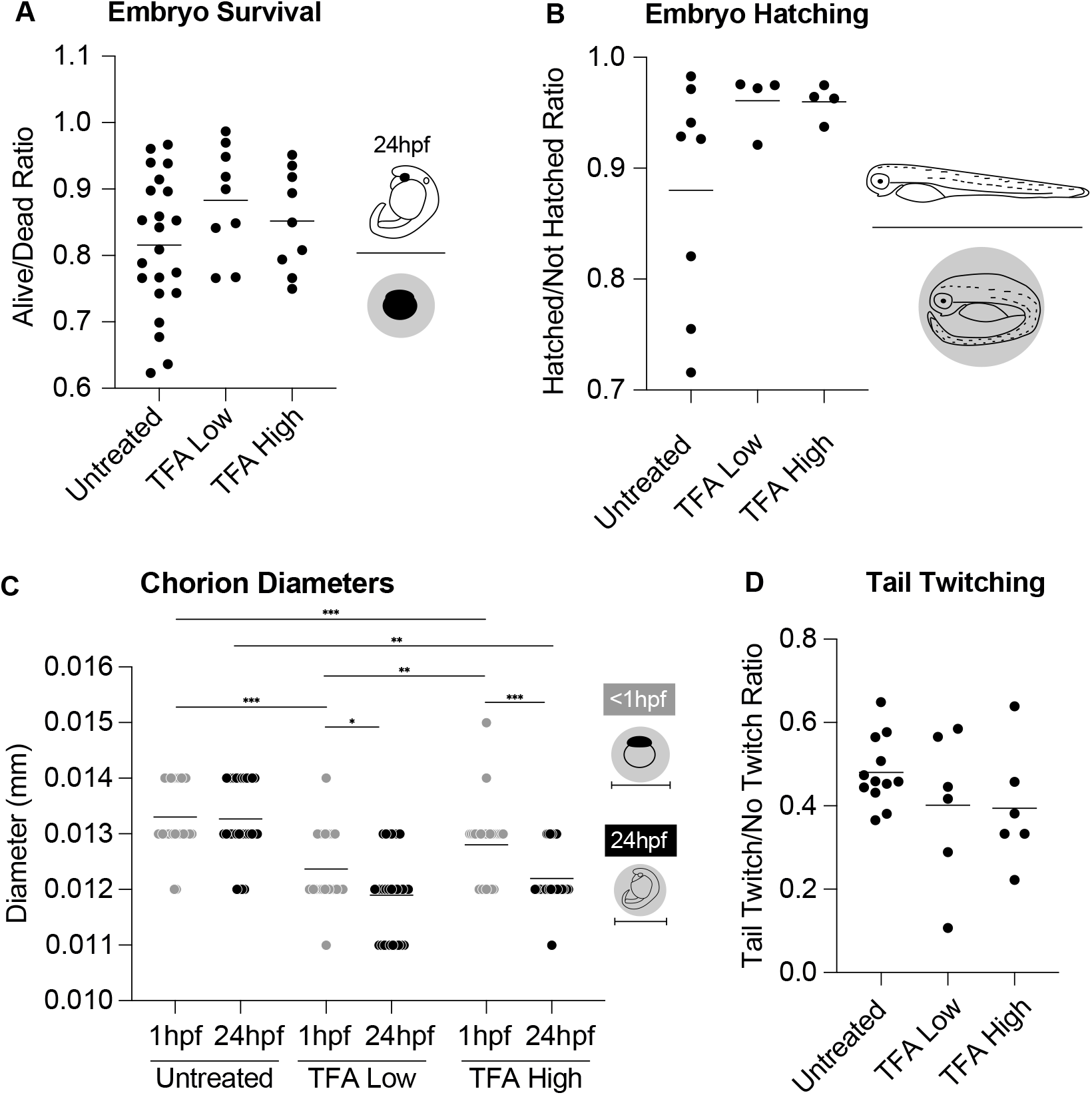
Developmental endpoints were assessed during early larval stages following TFA exposure initiated at the 1-cell stage; TFA low exposure: 45 ng/L and TFA high exposure: 1000 ng/L. (A) Jitter plot of embryo survival ratio at 24 hours post-fertilization (hpf) calculated as the number of alive over dead embryos, with a line representing the mean (B) Jitter plot for embryo hatching ratio at 72 hpf calculated with the number of hatched over not hatched embryos, with a line representing the mean hatching rate. (C) Jitter plot for average chorion diameter at 1- and 24 hpf, with a line for mean diameter. (D) Jitter plot for the tail twitch response at 24 hpf calculated with the number of embryos twitching over number not twitching in response to a manual poke. Significance was determined student paired t-test, p value depicted as * if < 0.05, ** if < 0.01, and *** if < 0.001.

Having established that TFA exposure does not compromise embryonic survival, we next examined whether early developmental timing was altered. Hatching rates were assessed at 72 hpf for each untreated and each exposure condition (Figure 1B). The ratio of hatched over unhatched embryos for both low and high TFA exposure groups exhibited an upward trend. To determine whether any changes in embryonic structure might underlie this trend of altered hatching phenotype, we next measured chorion diameter at early developmental time points. Chorion diameter measurements revealed a significant reduction within the first hour of being exposed to TFA for both exposure groups, with this reduction persisting and becoming more pronounced by 24 hpf (Figure 1C). Reduced chorion diameter may contribute to the observed acceleration in hatching, as TFA-exposed embryos experience increased spatial constraint within the chorion compared to untreated controls.

Given that early structural changes can sometimes precede functional alterations, we next assessed neuromuscular junction formation at 24 hpf using a tail twitch assay. Through the tail twitch assay, we observed no statistically significant difference within the TFA exposure groups compared to controls (Figure 1D). However, a consistent downward trend in response frequency was observed across both exposure groups. Although not statistically significant, this trend may indicate a delay in neuromuscular junction activity. Altogether, these early developmental changes motivated further exploration to determine if embryonic TFA exposure leads to persistent behavioral effects at later larval stages.

### Development impacts of short-chain PFAS persist causing anxiety-like behaviors in zebrafish larvae

Following early developmental impacts, we next assessed whether short-chain PFAS exposure produced functional consequences on larval behavior. Larval zebrafish at 5 dpf have a fully functional nervous system and stable free-swimming behavior which makes this stage ideal for detecting behavioral disruptions (Burgess & Burton, 2023; Colwill & Creton, 2011; Robertson et al., 2007), such as those caused by TFA exposure. Under normal conditions, larvae avoid light and display reduced movement in illuminated environments, whereas anxious larvae exhibit increased locomotor activity (Doszyn et al., 2024). We retested the same larvae were at 6 dpf to evaluate the persistence of behavioral changes (Figure 2A).

**Figure 2.**
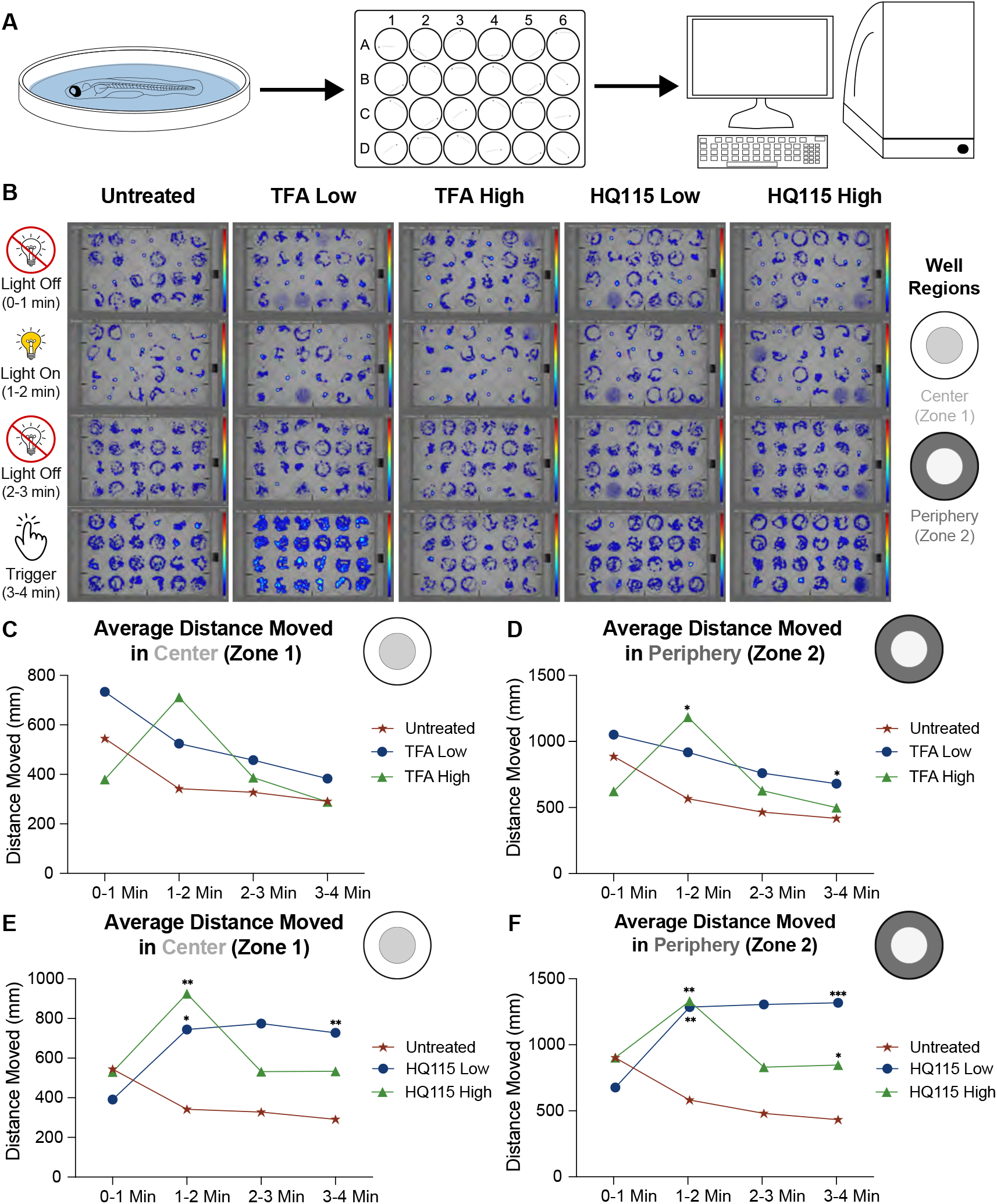
Recording of zebrafish larvae movement at 6 days post fertilization (dpf) following daily TFA and HQ115 exposure; TFA low exposure: 45 ng/L and TFA high exposure: 1000 ng/L and HQ115 low exposure: 400 ng/L and HQ115 high exposure: 8000 ng/L (A) Visual schematic to represent how each test was conducted. Larvae were placed into 24 well plate, 1 per well, and placed into the DanioVision chamber to undergo the predefined test of light off, light on, light off, and trigger stimulus. (B) Merged heatmap of the whole 24 well plate representing all larvae, separated into different time bins for each of the different predefined conditions. From left to right, the columns are untreated larvae, TFA low exposure, TFA high exposure, HQ115 low exposure, and HQ115 high exposure. From top to bottom, the rows are light off (0-1min), light on (1-2 min), light off (3-4 min), trigger stimulus (3-4 min), visually representing each time window for whether or not there was a light stimulus or a trigger stimulus which is a piston tapping the 24 well plate. The dark blue areas represent where the larvae spent most of their time and the high frequency circles represent the larvae frozen in that spot for an extended period. (C/E) Average distance moved in the center (Zone 1) of the well plate during each stimulus. (D/F) Average distance moved in the periphery (Zone 2) of the well plate during each stimulus. Significance was calculated using paired t-test comparing each exposed groups to the untreated group, p value depicted as * if < 0.05, ** if < 0.01, and *** if < 0.001.

Heatmap analyses revealed clear spatial differences between PFAS-exposed and control larvae (Figure 2B). TFA-exposed larvae move more and spent a larger proportion of time along the well walls, often swimming in circular patterns near the edge or remaining frozen against the wall.The high-density areas on the heatmaps indicate prolonged periods of inactivity or repeated wall-hugging behavior, suggesting elevated stress levels or anxiety-like responses.

This wall-hugging behavior, known as thigmotaxis, is commonly used as an indicator of anxiety in zebrafish (Schnörr et al., 2012). Larvae exposed daily to both low and high concentrations of TFA exhibited increased total movement and thigmotaxis (Figure 2). This behavior that persisted at 6 dpf and was also observed at 5 dpf, with similar trends (Supp Figure 2). Additionally, the TFA exposed larvae displayed hyperactivity followed by reduced movement or freezing toward the end, suggesting fatigue or a stress response (Figure 2C and 2D). The timing of peak hyperactivity periods were stimuli dependent. TFA high exposure group peak activity was during the light stimulus (1-2 min), whereas TFA low exposure group peak activity was during the startle stimulus (3-4 min). This hyperactivity, specifically during the light or trigger stimulus, indicates an anxiety-like response. When comparing the overall movements across zones, center (Zone 1) and periphery (Zone 2), larvae exposed to both concentrations of TFA had increased movement in the periphery of the well (Zone 2). This is consistent with enhanced thigmotaxis and is indicative of higher anxiety response compared to untreated controls.

To determine whether these behavioral effects were unique to TFA or shared across short-chain PFAS, larvae were also exposed to Bis(trifluoromethane)sulfonimide lithium salt (HQ115), another short-chain PFAS, at environmentally relevant low and high concentrations (Sands, 2024). HQ115-exposed larvae exhibited a behavioral profile similar to TFA-exposed larvae (Supp figure 2 and 4). Repeating the test at 6 dpf confirmed that these effects persisted for both TFA and HQ115, with larvae exposed to HQ115 showing increased movement and hyperactivity in response to both light and startle stimuli, suggesting that both short-chain PFAS induce anxiety-like behavior (Figure 2E and 2F).

Together, these results suggest that early exposure to short-chain PFAS produces lasting anxiety-like behavior in developing zebrafish. The similar responses in larvae observed for TFA and HQ115 indicate that hyperactivity and thigmotaxis may be common features of PFAS exposure, reflecting persistent neural alterations during development.

### Short-chain PFAS alter neuromuscular and stress-related protein expression during critical windows of zebrafish development

To explore potential molecular mechanisms underlying the behavioral changes observed in larvae, we examined protein expression in zebrafish embryos exposed to TFA at 24 and 48 hpf. These developmental timepoints are critical windows for neuromuscular development, organogenesis, and nervous system formation (Liu et al., 2023; Panzer et al., 2005), and can be particularly sensitive to toxicant exposure. Proteomic analysis revealed significant dosage and time dependent alterations in protein expression following TFA exposure, giving insight into early molecular disruptions that could contribute to later behavioral outcomes.

At 24 hpf, embryos exposed to both low and high concentrations of TFA have significant alternations in protein expression compared to untreated controls (Figures 3A and 3B, Supp Table 1 and Supp Table 2). Differentially expressed proteins (Table1A and Table2A) were involved in pathways related to muscle activity, neuronal signaling, and stress response, and have been implicated in several human diseases (Table 1B and Table 2B). In the high concentration group, proteins Pvalb2 (Q9I8V0), Hyou1 (Q7ZUW2), and Actbb (Q7ZVF9) were upregulated. These proteins are involved in the processes of muscle relaxation, swimming performance, and anxiety regulation (Xu et al., 2000), indicating early activation of neuromuscular and stress-related pathways. In the low concentration group, the upregulated proteins along with Actbb (Q7ZVF9) were also Krt8 (Q6NWF6), and Nme3 (Q9PTF3), both of which are involved in swim endurance, oxidative stress pathway, and energy metabolism (Abu-Taha et al., 2017; Song et al., 2004). These molecular changes from both exposure groups are consistent with the hyperactivity and elevated stress responses observed during later larval behavioral assays. Additionally, both TFA exposure groups at 24 hpf exhibited downregulation of Mybb1a (Q6DRL5), a protein involved in circadian rhythm regulation (Hara et al., 2009). Disruption of circadian pathways during early development has been linked to altered stress responsiveness and anxiety-like behaviors (Pintos, 2023; Yang, 2018). Together, these early protein changes provide a molecular explanation for the hyperactivity and anxiety-like behaviors observed in the behavioral assays.

**Table 1.**
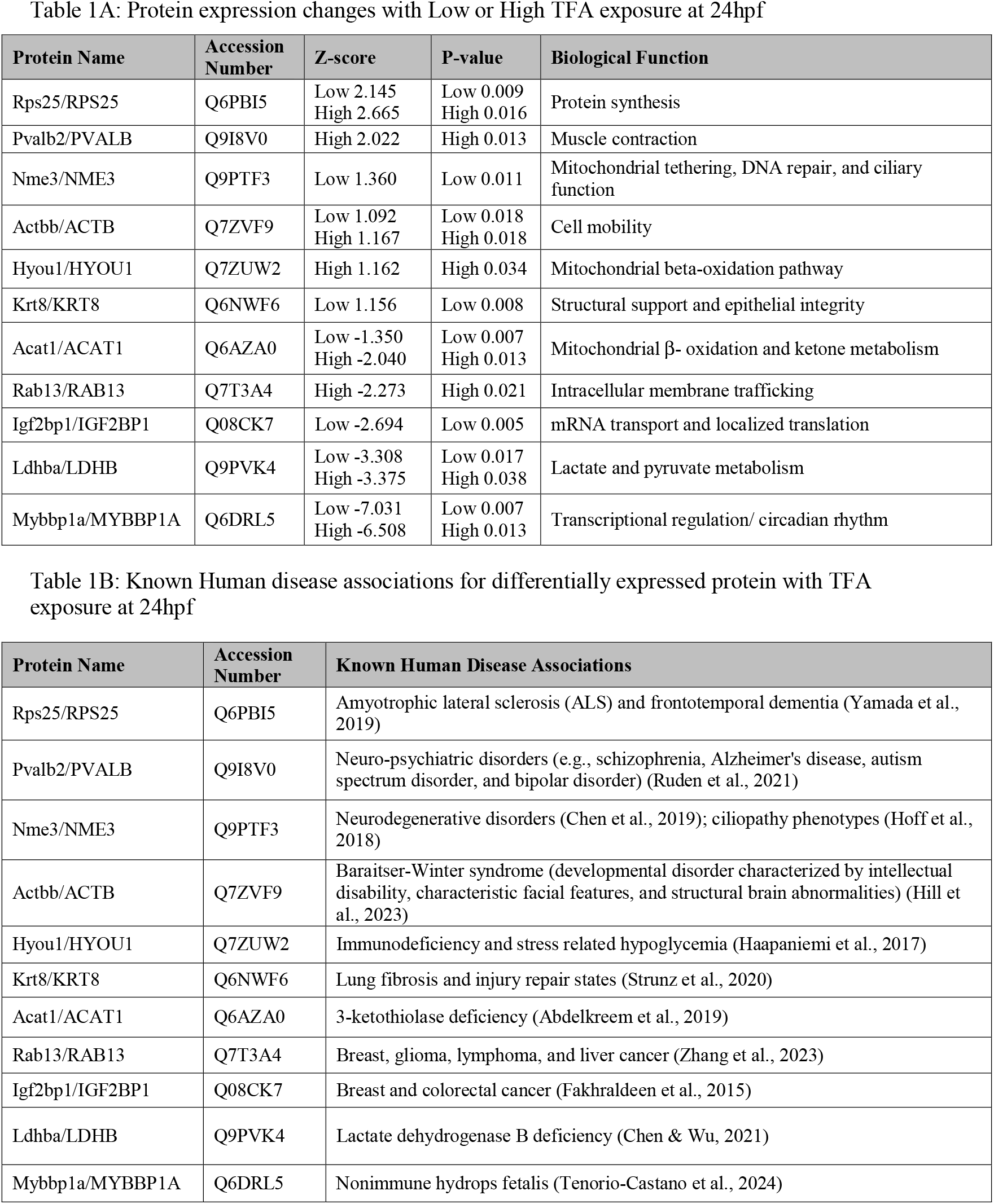
(A) Top differentially expressed proteins following TFA exposure at 24 hpf compared to untreated controls. Low TFA exposure: 45 ng/L; High TFA exposure: 1000 ng/L. Protein names are listed for zebrafish and their human equivalents (Zebrafish/Human). Z-scores (positive = upregulated; negative = downregulated), p-values, and protein function. Protein names and biological functions were obtained from UniProt and the Human Protein Atlas. (B) For each differentially expressed protein, known associated human diseases are provided.

**Table 2.**
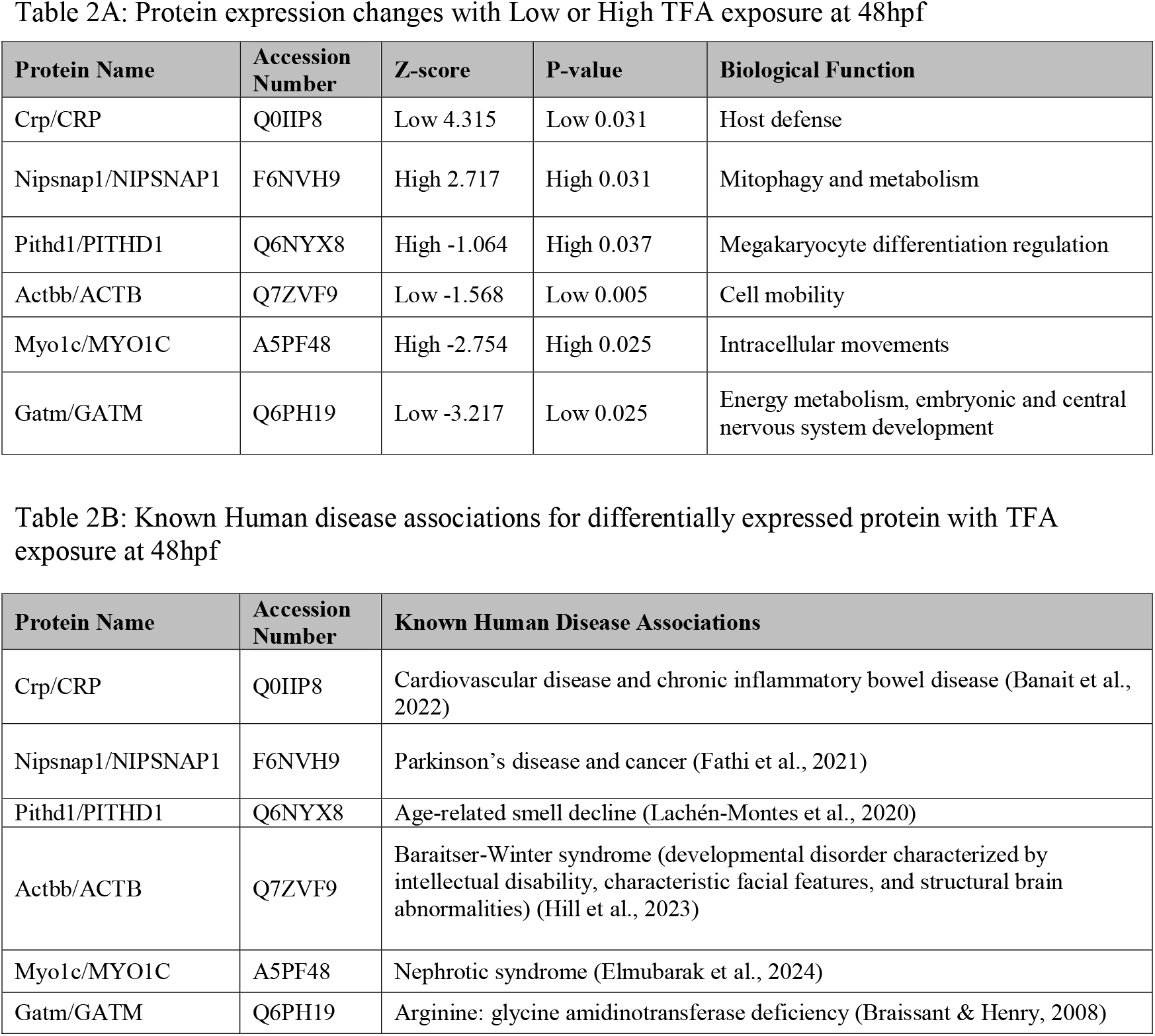
(A) Top differentially expressed proteins following TFA exposure at 48 hpf compared to untreated controls. Low TFA exposure: 45 ng/L; High TFA exposure: 1000 ng/L. Protein names are listed for zebrafish and their human equivalents (Zebrafish/Human). Z-scores (positive = upregulated; negative = downregulated), p-values, and protein function. Protein names and biological functions were obtained from UniProt and the Human Protein Atlas. (B) For each differentially expressed protein, known associated human diseases are provided.

**Figure 3.**
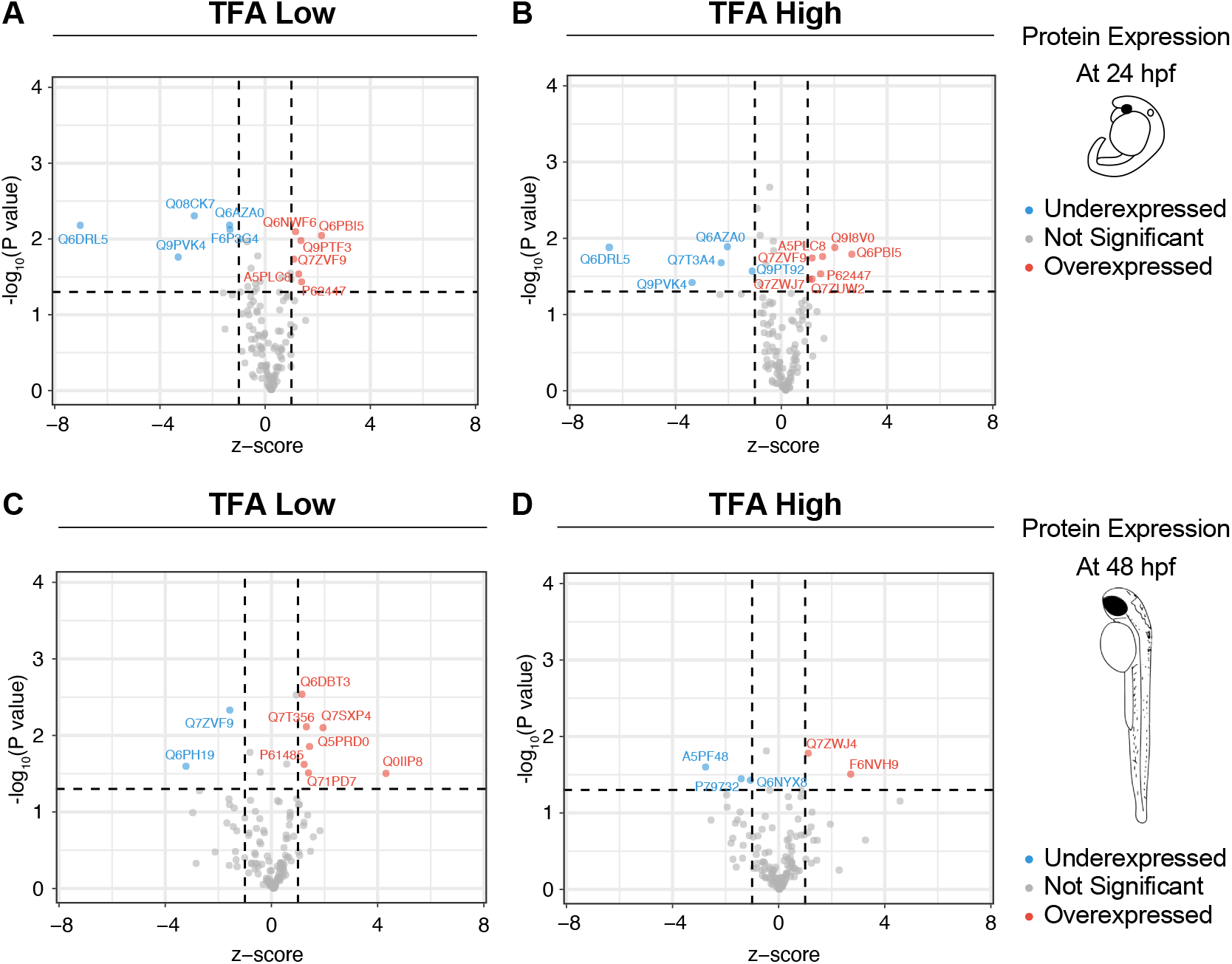
Volcano plot depicting differential protein expression for 24 and 48 hpf TFA exposed zebrafish embryos compared to untreated controls. All embryos were exposed to their respective TFA concentration at 1 cell then were collected at 24 hpf or 48 hpf to test protein expression. Top panel represents the embryos exposed for 24 hpf with (A) being TFA low (45 ng/L) exposed embryos and (B) being TFA high (1000 ng/L) exposed embryos. Bottom panel represents the embryos exposed for 48 hpf with (C) being TFA low (45 ng/L) embryos and (D) being TFA high (1000 ng/L) embryos. Red dots indicate proteins significantly upregulated compared to control, and blue dots indicate proteins significantly downregulated compared to controls.

By 48 hpf, protein expression profiles shifted toward predominantly downregulation, particularly in the high TFA exposure group (Figure 3D), reflecting continued developmental disruptions. Proteins Actbb (Q7ZVF9), Myo1c (A5PF48), and Pithd1 (Q6NYX8) which are involved in swim behavior, muscle function, fin development, and stress circuits (Arif et al., 2013; Sohn et al., 2009), were downregulated. This suggests impaired swim and muscle endurance, and neuromotor capacity, consistent with the observed decline in swim activity following initial hyperactivity during behavioral testing for the high exposure group. In the low concentration, downregulation of Gatm (Q6PH19), which is associated with muscle strength and fatigue (Wang et al., 2007), was observed. This is indicative of diminished capacity for sustained movement which is consistent with the gradual decrease in activity during behavioral assays. During behavioral assays, this low concentration group, rarely had peaks of extremely high movement, and even after small bouts of increased movement the total distance the larvae moved continued to decrease.

To determine whether these molecular changes are unique to TFA or shared with other environmentally relevant but long chain PFAS, we performed proteomic analysis on embryos exposed to perfluorooctanoic acid (PFOA) at the same time points (Figure 4). Three proteins, Acat1 (Q6AZA0), Pvalb2 (Q9I8V0), and Igf2bp1 (Q08CK7), were affected by both PFAS, having overlapping roles in swimming-related processes, including muscle relaxation, energy metabolism, and startle response (Gu et al., 2020; Xu et al., 2000). Notably, the timing of protein expression changes differed slightly, such that for PFOA, at 48 hpf, the low-dose group showed downregulation of Pvalb2 (Q9I8V0) and Igf2bp1 (Q08CK7), while the high-dose group at 24 hpf exhibited downregulation of Acat1 (Q6AZA0). In contrast to PFOA, with TFA exposure, all three proteins were already altered at 24 hpf, with Pvalb2 (Q9I8V0) being the only protein that was over-expressed. All these proteins are involved with some aspect of how the fish swim whether it is muscle relaxation, energy, or startle response. This similarity suggests that short-chain PFAS, such as TFA, can induce molecular disruptions comparable to those caused by longer-chain PFAS like PFOA (Supplementary Figure 6), and that some of these effects even occur at an earlier developmental time point.

Notably, several of the differential expressed proteins are implicated in processes critical for early embryonic development, including ciliogenesis, mRNA localization, epithelial integrity, and cellular metabolism, and are linked to human neurodevelopmental and metabolic disorders (Table 1 and 2). Collectively, our findings demonstrate that TFA exposure impacts key proteins involved in muscle function, energy metabolism, and neural formation in a dose- and time-dependence manner. Early overexpression of proteins at 24 hpf may drive the hyperactivity and anxiety-like behaviors observed in larvae, while subsequent downregulation by 48 hpf likely contributes to the reduced muscle endurance and neuromuscular performance. The overlap of protein expression with PFOA-induced changes further highlights that short-chain PFAS can produce molecular effects akin to long-chain PFAS. Overall, these results support the conclusion that TFA negatively influences early developmental processes, with higher concentrations producing greater effects, but lower concentrations still causing measurable molecular and behavioral disruptions.

**Figure 4.**
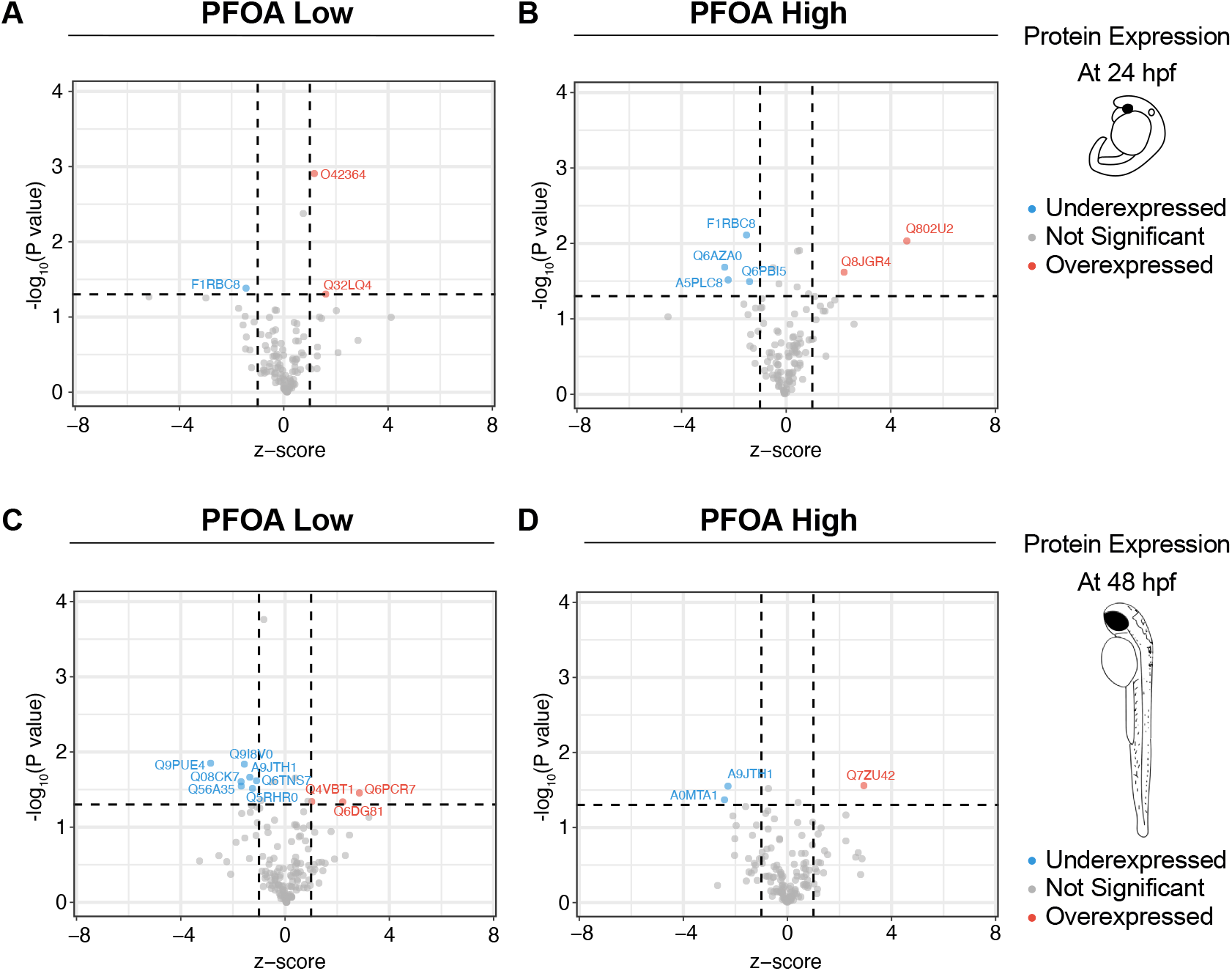
Volcano plot depicting differential protein expression for 24 and 48 hpf PFOA exposed zebrafish embryos compared to untreated controls. All embryos were exposed to their respective PFOA concentration at 1 cell then were collected at 24 hpf or 48 hpf to test protein expression. Top panel represents the embryos exposed for 24 hpf with (A) being PFOA low (700 ng/L) exposed embryos and (B) being PFOA high (10,400,000 ng/L) exposed embryos. Bottom panel represents the embryos exposed for 48 hpf with (C) being PFOA low (700 ng/L) embryos and (D) being PFOA high (10,400,000 ng/L) embryos. Red dots indicate proteins significantly upregulated compared to control, and blue dots indicate proteins significantly downregulated compared to controls.

## Conclusion

This study demonstrated that early-life exposure to the short chain PFAS TFA, at both low and high dosages, produces measurable developmental, behavioral, and protein expression alternations in zebrafish. Although TFA exposure did not significantly affect embryo mortality at either low and high dosages, exposed embryos exhibited reduced chorion diameter as early as 1 hpf and continuing at 24 hpf. Furthermore, at 24 hpf embryos exhibited a downward trend for tail twitching indicating disruptions to neuromuscular junction formation. These results suggested that early exposure to TFA may disrupt fundamental developmental processes which warranted further investigation into their swim behavior at a later developmental timepoint.

Behavioral assays conducted at 5 and 6 dpf revealed increased locomotor activity and heightened thigmotaxis compared to untreated controls (Figure 5A and Supp Figure 3). These effects were stimulus-dependent and consistent with anxiety-like behavior, suggesting that early TFA exposure can have lasting neurobehavioral consequences. Together the developmental and behavioral data suggests that TFA exposure disrupts neuromuscular developmental processes that are critical for normal behavioral responses.

**Figure 5.**
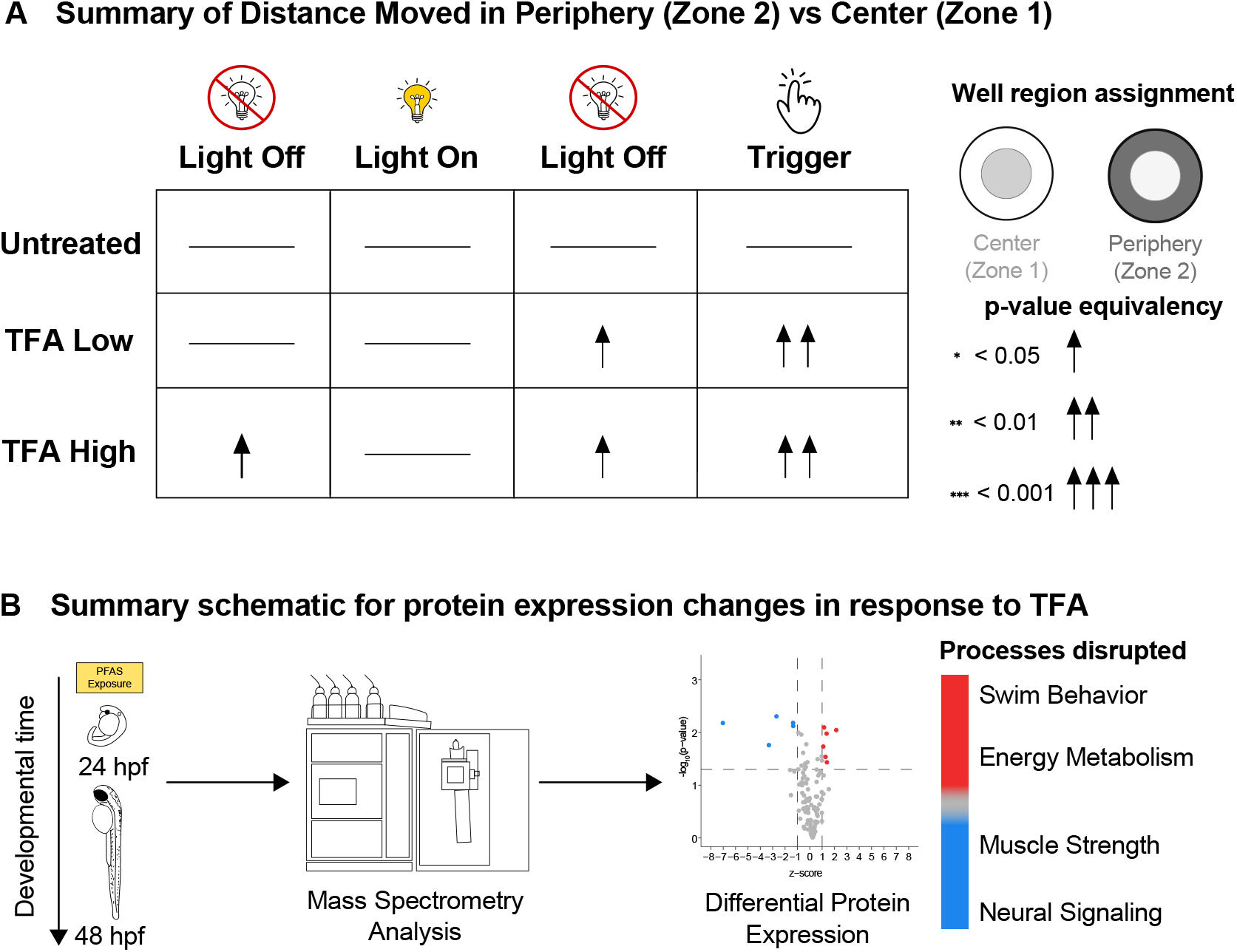
(A) Schematic illustrating larval preference for the periphery over the center. Behavioral measures represent the most significantly altered responses following short-chain PFAS exposure. Straight lines denote untreated controls used for comparison. Upward-facing arrows indicate increased distance traveled relative to controls, and arrow thickness reflects the degree of statistical significance. (B) Schematic depicting time- and dose-dependent protein expression changes in response to short-chain PFAS exposure, TFA and HQ115. Disrupted biological processes inferred from differential protein expression are listed on the right.

To explore the molecular mechanisms underlying these developmental and behavioral effects, we then conducted proteomic profiling. This analysis revealed dose- and time-dependent disruptions in proteins involved in swim behavior, muscle strength, and neural signaling pathways related to stress and anxiety (Figure 5B). Early upregulation of proteins associated with swimming performance and stress responses at 24 hpf was followed by downregulation of proteins needed for muscle endurance and neuromotor stability at 48 hpf. Importantly, overlapping protein expression changes between TFA and PFOA indicate that TFA can engage similar molecular pathways. These results provide a mechanistic link between early-stage protein disruptions and the functional deficits observed at later larval stages with embryonic exposure of PFAS.

Collectively, these findings demonstrate that the short chain PFAS TFA may pose underestimated health risks, challenging the assumption that TFA is an inherently safer alternative. Instead, our data suggest that TFA exposure during early development can produce lasting molecular and behavioral effects, highlighting the need for further evaluation of the short chain PFAS toxicity.

## Acknowledgments

This research was supported by RCMI grant U5MD012392 awarded to DK and the RCMI pilot grant awarded to ZA, and the NC PFAS Collaboratory grant for Molecular mechanisms of PFAS. We want to also acknowledge the Julius L. Chambers Biomedical/Biotechnology Research Institute (BBRI) at North Carolina Central University for providing research facilities and resources, and thank all core faculty and staff, including Dr. Derek Norford. We want to also thank the high school students from the SRSP program, Minchae Joo and Christian Niu. Additionally, would like to thank the Georgetown Summer PACT Program, Dr. Jan Blancato, and Dr. Caleb McKinney for supporting undergraduate research experience and professional development. Finally, we would like to also thank Leslimar Rios-Colon for her support at RTI International.

## Figure legends

**Supp Figure 1** Developmental endpoints were assessed during early larval stages following TFA exposure initiated at the 1-cell stage; HQ115 low exposure: 400 ng/L and HQ115 high exposure: 8000 ng/L. (A) Jitter plot of embryo survival ratio at 24 hours post-fertilization (hpf) calculated as the number of alive over dead embryos, with a line representing the mean (B) Jitter plot for embryo hatching ratio at 72 hpf calculated with the number of hatched over not hatched embryos, with a line representing the mean hatching rate. (C) Jitter plot for average chorion diameter at 1- and 24 hpf, with a line for mean diameter. (D) Jitter plot for the tail twitch response at 24 hpf calculated with the number of embryos twitching over number not twitching in response to a manual poke. Significance was determined student paired t-test, p value depicted as * if < 0.05, ** if < 0.01, and *** if < 0.001.

**Supp Figure 2** Recording of zebrafish larvae movement at 5 days post fertilization (dpf) following daily TFA and HQ115 exposure; TFA low exposure: 45 ng/L and TFA high exposure: 1000 ng/L and HQ115 low exposure: 400 ng/L and HQ115 high exposure: 8000 ng/L (A) Visual schematic to represent how each test was conducted. Larvae were placed into 24 well plate, 1 per well, and placed into the DanioVision chamber to undergo the predefined test of light off, light on, light off, and trigger stimulus. (B) Merged heatmap of the whole 24 well plate representing all larvae, separated into different time bins for each of the different predefined conditions. From left to right, the columns are untreated larvae, TFA low exposure, TFA high exposure, HQ115 low exposure, and HQ115 high exposure. From top to bottom, the rows are light off (0-1min), light on (1-2 min), light off (3-4 min), trigger stimulus (3-4 min), visually representing each time window for whether or not there was a light stimulus or a trigger stimulus which is a piston tapping the 24 well plate. The dark blue areas represent where the larvae spent most of their time and the high frequency circles represent the larvae frozen in that spot for an extended period. (C/E) Average distance moved in the center (Zone 1) of the well plate during each stimulus. (D/F) Average distance moved in the periphery (Zone 2) of the well plate during each stimulus. Significance was calculated using paired t-test comparing each exposed groups to the untreated group, p value depicted as * if < 0.05, ** if < 0.01, and *** if < 0.001.

**Supp Figure 3** Recording of zebrafish larvae movement at 6 days post fertilization (dpf) following daily TFA; TFA low exposure: 45 ng/L and TFA high exposure: 1000 ng/L. From left to right, the y-axis represents light off (0-1min), light on (1-2 min), light off (3-4 min), trigger stimulus (3-4 min). (A) Average velocity moved in the center (Zone 1) of the well plate during each stimulus. (B) Average velocity moved in the periphery (Zone 2) of the well plate during each stimulus. Significance was calculated using paired t-test comparing each exposed groups to the untreated group, p-value depicted as * if < 0.05, ** if < 0.01, and *** if < 0.001.

**Supp Figure 4** Recording of zebrafish larvae movement at 6 days post fertilization (dpf) following daily HQ115; HQ115 low exposure: 400 ng/L and HQ115 high exposure: 8000 ng/L. From left to right, the y-axis represents light off (0-1min), light on (1-2 min), light off (3-4 min), trigger stimulus (3-4 min). (A) Average velocity moved in the center (Zone 1) of the well plate during each stimulus. (B) Average velocity moved in the periphery (Zone 2) of the well plate during each stimulus. Significance was calculated using paired t-test comparing each exposed groups to the untreated group, p-value depicted as * if < 0.05, ** if < 0.01, and *** if < 0.001.

**Supp Figure 5** Schematic representation of larval preference for the center and the periphery compared to untreated larvae. Behavioral measures shown reflect the most significantly altered responses following short-chain PFAS exposure. Straight lines denote untreated controls used for comparison. Upward-facing arrows indicate increased distance moved relative to untreated embryos, while arrow thickness corresponds to the degree of statistical significance.

**Supp Figure 6** UpSet plot depicting the intersection of differentially expressed proteins across exposure conditions relative to untreated controls. Bars represent the number of proteins shared between one or more treatment groups, with connected points indicating specific condition overlaps. The x-axis denotes the exposure conditions included in each comparison. A) For all samples collected 24 hpf. B) For all samples collected at 48 hpf. TFA low exposure: 45 ng/L and TFA high exposure: 1000 ng/L. PFOA low exposure: 400 ng/L and PFOA high exposure: 10,400,000 ng/L.

