## Supplementary material for "Short-chain PFAS exposure alters embryonic development and behavior in zebrafish": Supp Figure: SuppFigures_compressed.pdf

**Supp Figure 1**

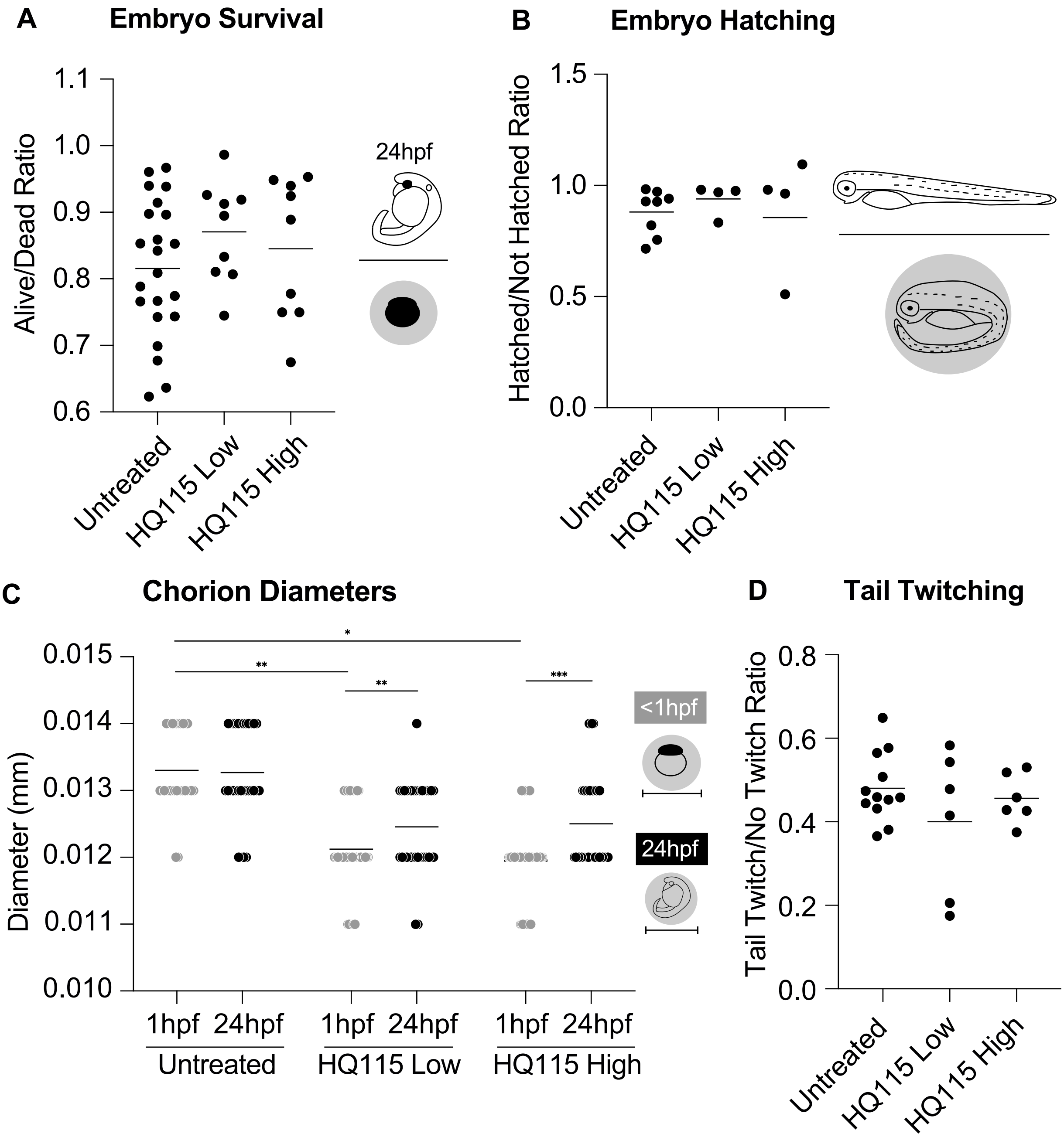


**Supp Figure 1** Developmental endpoints were assessed during early larval stages following TFA exposure initiated at the 1-cell stage; HQ115 low exposure: 400 ng/L and HQ115 high exposure: 8000 ng/L. (A) Jitter plot of embryo survival ratio at 24 hours post-fertilization (hpf) calculated as the number of alive over dead embryos, with a line representing the mean (B) Jitter plot for embryo hatching ratio at 72 hpf calculated with the number of hatched over not hatched embryos, with a line representing the mean hatching rate. (C) Jitter plot for average chorion diameter at 1- and 24 hpf, with a line for mean diameter. (D) Jitter plot for the tail twitch response at 24 hpf calculated with the number of embryos twitching over number not twitching in response to a manual poke. Significance was determined student paired t-test, p value depicted as * if $< 0.05$, ** if $< 0.01$, and *** if $< 0.001$.

Supp Figure 2

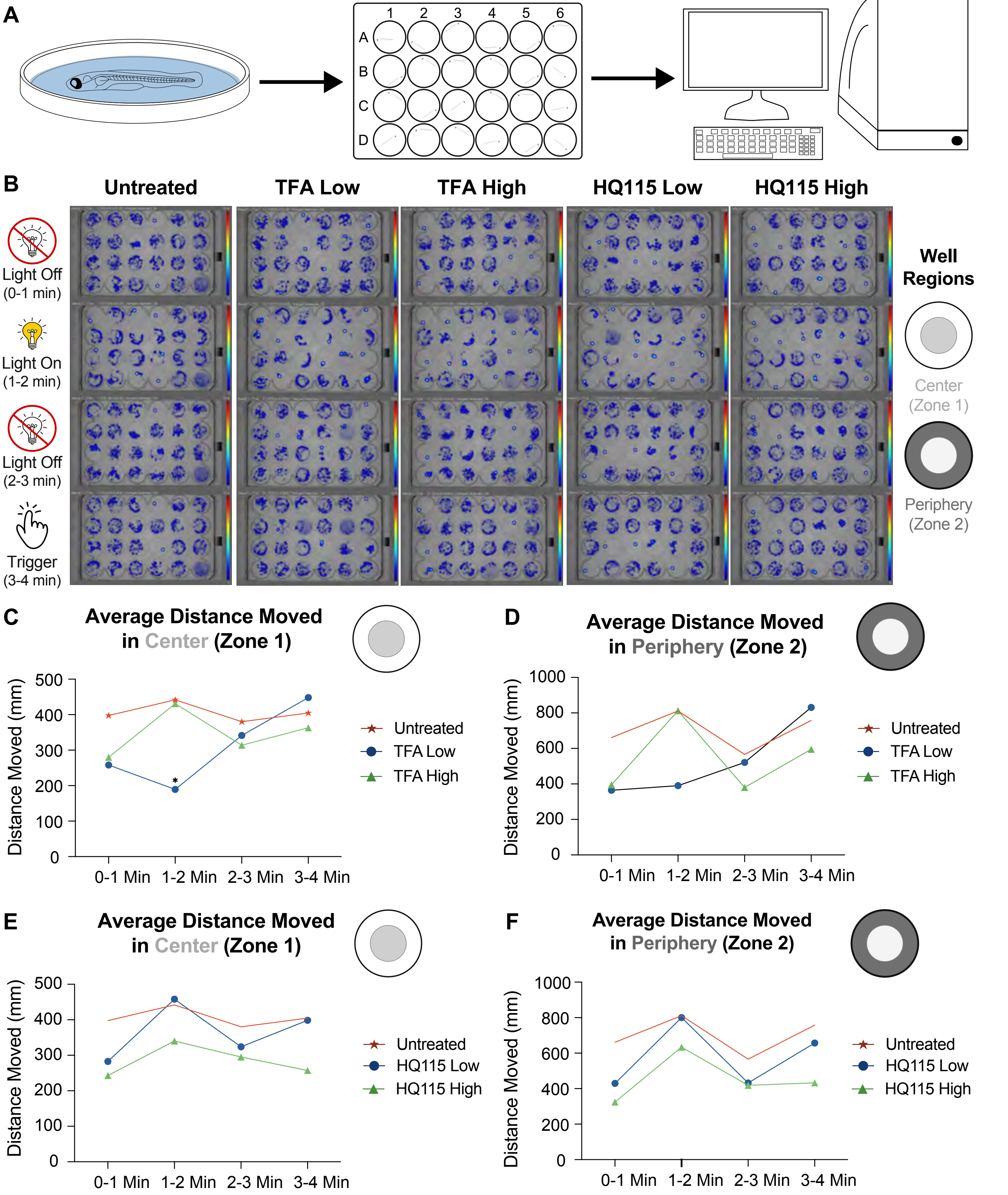


**Supp Figure 2** Recording of zebrafish larvae movement at **5 days post fertilization (dpf)** following daily TFA and HQ115 exposure; TFA low exposure: 45 ng/L and TFA high exposure: 1000 ng/L and HQ115 low exposure: 400 ng/L and HQ115 high exposure: 8000 ng/L (A) Visual schematic to represent how each test was conducted. Larvae were placed into 24 well plate, 1 per well, and placed into the DanioVision chamber to undergo the predefined test of light off, light on, light off, and trigger stimulus. (B) Merged heatmap of the whole 24 well plate representing all larvae, separated into different time bins for each of the different predefined conditions. From left to right, the columns are untreated larvae, TFA low exposure, TFA high exposure, HQ115 low exposure, and HQ115 high exposure. From top to bottom, the rows are light off (0-1min), light on (1-2 min), light off (3-4 min), trigger stimulus (3-4 min), visually representing each time window for whether or not there was a light stimulus or a trigger stimulus which is a piston tapping the 24 well plate. The dark blue areas represent where the larvae spent most of their time and the high frequency circles represent the larvae frozen in that spot for an extended period. (C/E) Average distance moved in the center (Zone 1) of the well plate during each stimulus. (D/F) Average distance moved in the periphery (Zone 2) of the well plate during each stimulus. Significance was calculated using paired t-test comparing each exposed groups to the untreated group, p value depicted as * if < 0.05, ** if < 0.01, and *** if < 0.001.

**Supp Figure 3**

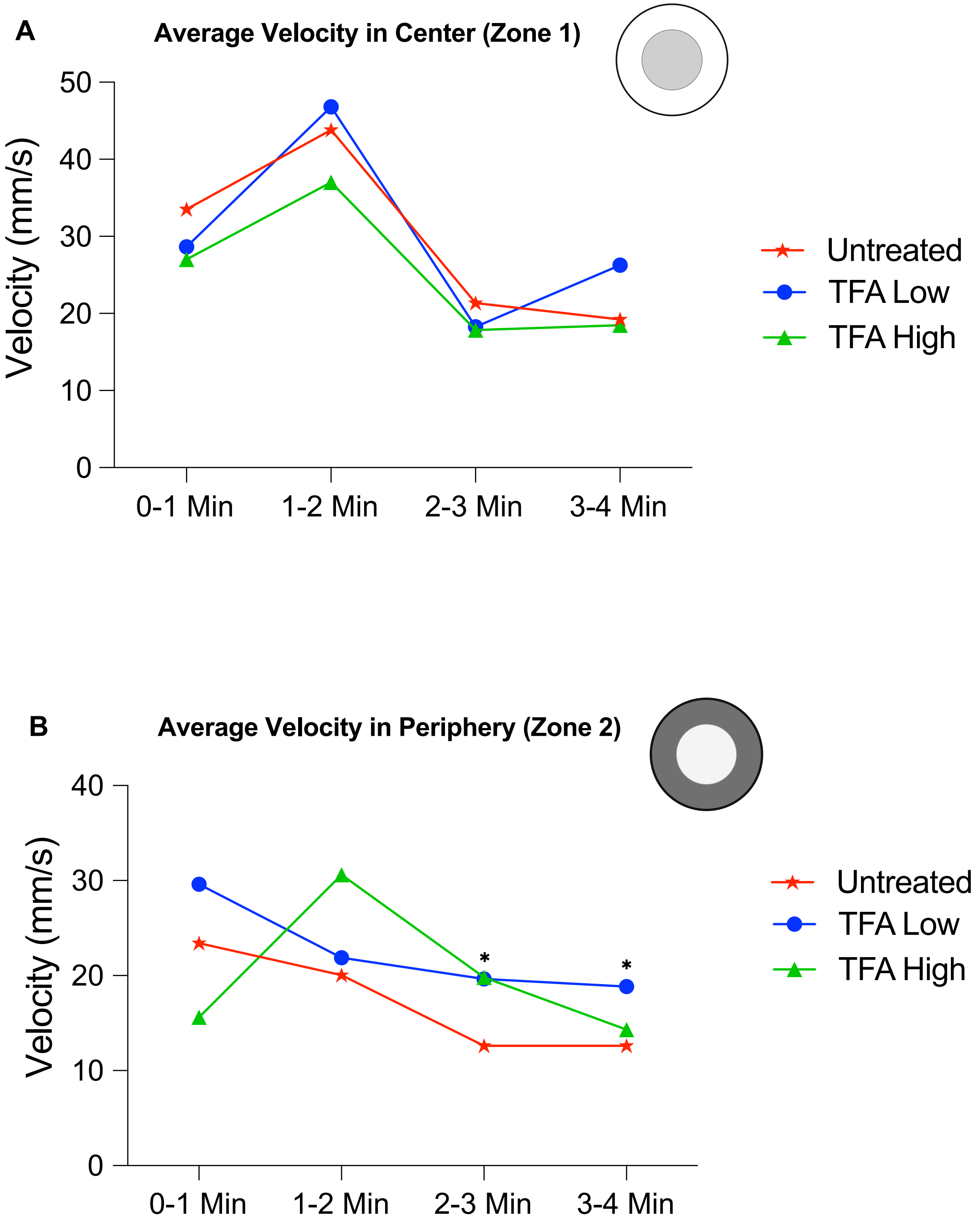


**Supp Figure 3** Recording of zebrafish larvae movement at 6 days post fertilization (dpf) following daily TFA; TFA low exposure: 45 ng/L and TFA high exposure: 1000 ng/L. From left to right, the y-axis represents light off (0-1min), light on (1-2 min), light off (3-4 min), trigger stimulus (3-4 min). (A) Average velocity moved in the center (Zone 1) of the well plate during each stimulus. (B) Average velocity moved in the periphery (Zone 2) of the well plate during each stimulus. Significance was calculated using paired t-test comparing each exposed groups to the untreated group, p-value depicted as * if < 0.05, ** if < 0.01, and *** if < 0.001.

**Supp Figure 4**

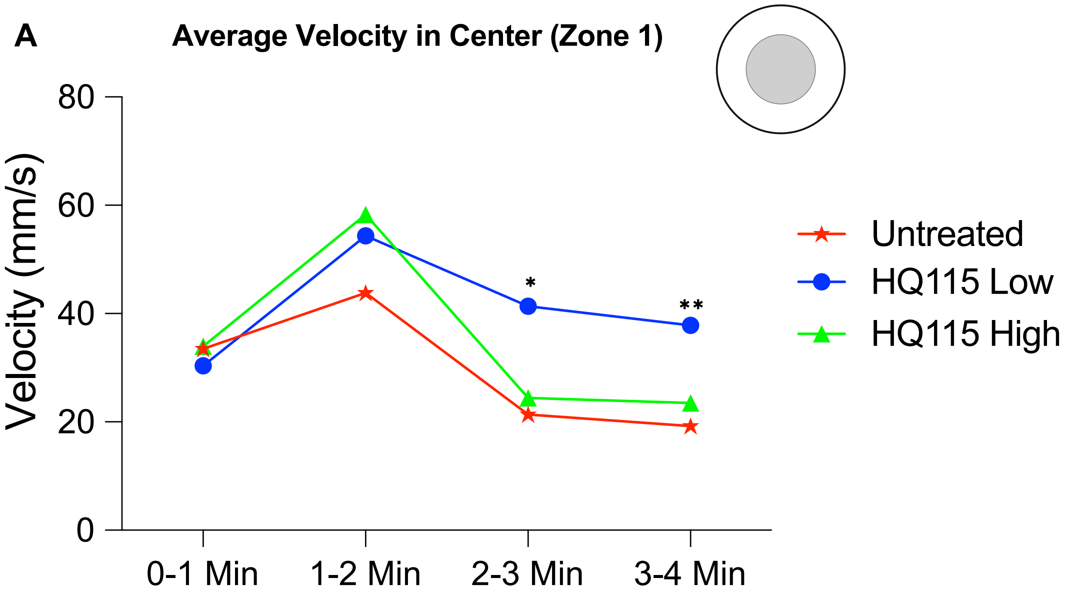


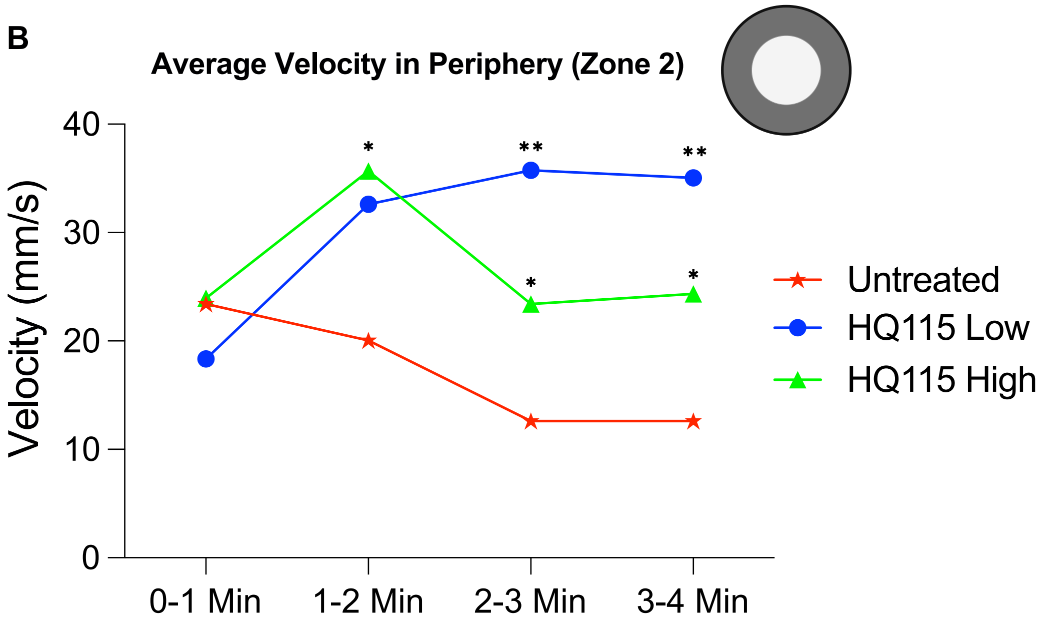


**Supp Figure 4** Recording of zebrafish larvae movement at 6 days post fertilization (dpf) following daily HQ115; HQ115 low exposure: 400 ng/L and HQ115 high exposure: 8000 ng/L. From left to right, the y-axis represents light off (0-1min), light on (1-2 min), light off (3-4 min), trigger stimulus (3-4 min). (A) Average velocity moved in the center (Zone 1) of the well plate during each stimulus. (B) Average velocity moved in the periphery (Zone 2) of the well plate during each stimulus. Significance was calculated using paired t-test comparing each exposed groups to the untreated group, p-value depicted as * if < 0.05, ** if < 0.01, and *** if  < 0.001.

### Supp Figure 5

#### Summary of Distance Moved in Center (Zone 1) Compared to Untreated

| | Light Off | Light On | Light Off | Trigger |
|---|---|---|---|---|
| TFA Low | ——— | ——— | ——— | ——— |
| TFA High | ——— | ↑ | ——— | ——— |
| HQ115 Low | ——— | ↑ | ↑ | ↑↑ |
| HQ115 High | ——— | ↑↑ | ——— | ——— |

**Well region**

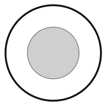


**p-value equivalency**

- * < 0.05 ↑
- ** < 0.01 ↑↑
- *** < 0.001 ↑↑↑

#### Summary of Distance Moved in Periphery (Zone 2) Compared to Untreated

| | Light Off | Light On | Light Off | Trigger |
|---|---|---|---|---|
| TFA Low | ——— | ——— | ——— | ↑ |
| TFA High | ——— | ↑ | ——— | ——— |
| HQ115 Low | ——— | ↑↑ | ↑↑ | ↑↑↑ |
| HQ115 High | ——— | ↑↑ | ——— | ↑ |

**Well region**

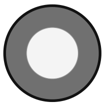


**p-value equivalency**

- * < 0.05 ↑
- ** < 0.01 ↑↑
- *** < 0.001 ↑↑↑

**Supp Figure 5** Schematic representation of larval preference for the center and the periphery compared to untreated larvae. Behavioral measures shown reflect the most significantly altered responses following short-chain PFAS exposure. Straight lines denote untreated controls used for comparison. Upward-facing arrows indicate increased distance moved relative to untreated embryos, while arrow thickness corresponds to the degree of statistical significance.

**Supp Figure 6**

**A** **Common differentially expressed proteins across treatment groups relative to untreated controls at 24 hpf**

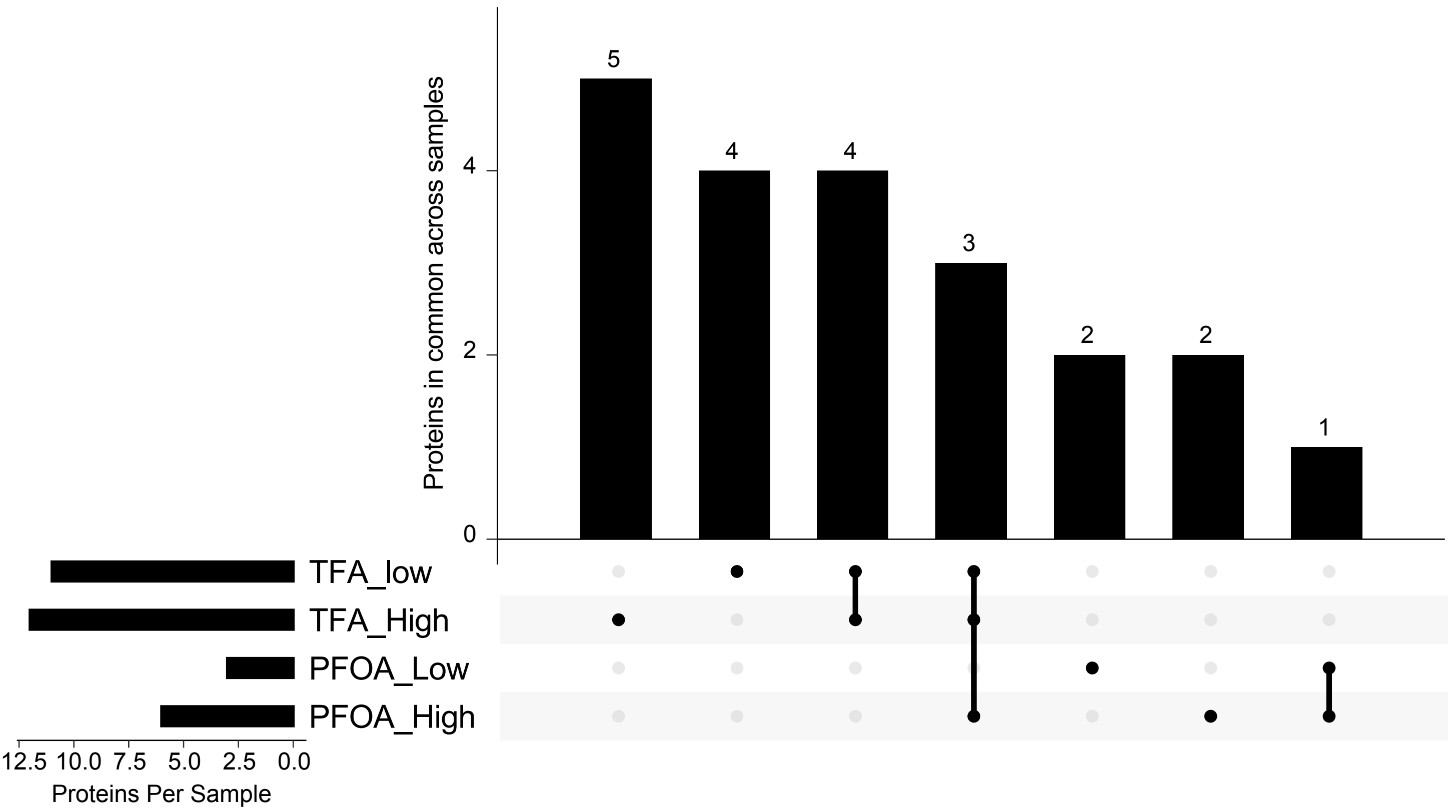


**B** **Common differentially expressed proteins across treatment groups relative to untreated controls at 48 hpf**

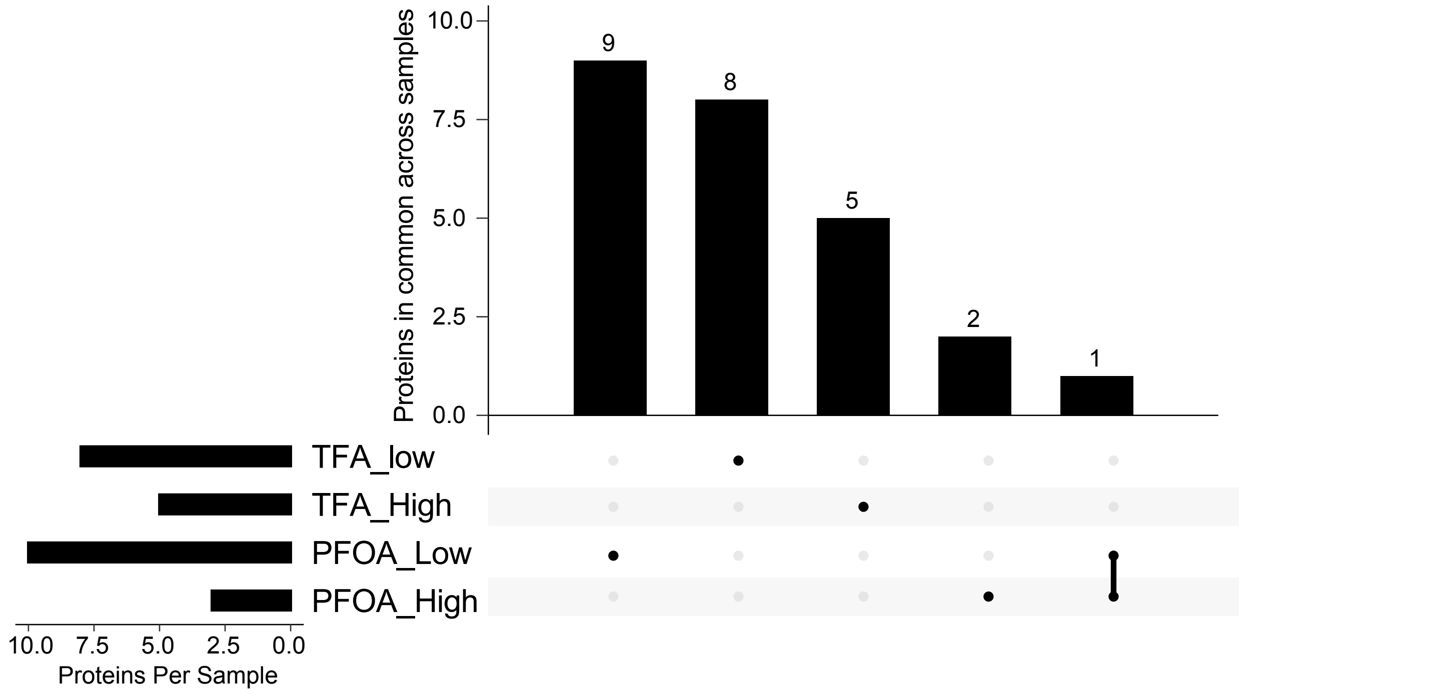


**Supp Figure 6** UpSet plot depicting the intersection of differentially expressed proteins across exposure conditions relative to untreated controls. Bars represent the number of proteins shared between one or more treatment groups, with connected points indicating specific condition overlaps. The x-axis denotes the exposure conditions included in each comparison.
A) For all samples collected 24 hpf. B) For all samples collected at 48 hpf. TFA low exposure: 45 ng/L and TFA high exposure: 1000 ng/L. PFOA low exposure: 400 ng/L and PFOA high exposure: 10,400,000 ng/L.
